# Replicate Engineered Virtual Patient Populations as Surrogates for Real Patient-Level Data

**DOI:** 10.1101/308403

**Authors:** Francis J. Alenghat

## Abstract

**Objectives:** To demonstrate a new method for generating virtual, individual-level data by testing it on a known clinical trial population.

**Design:** Virtualization of aggregate data from a clinical trial.

**Setting:** Virtual

**Participants:** 936,100 virtual patients

**Interventions:** None

**Main Outcomes Measures:** Odds ratios for adverse outcomes in virtual patient populations compared to clinical trial participants.

**Methods:** The replicate engineered virtual patient populations (RE-ViPPs) method, based on aggregate cross-tabulated categorical population data, does not require access to individual-level data. Using sequential regression combined with randomization, it generates virtual individual patients to comprise populations that, on average, closely resemble the real population in question. The method is validated by applying it to aggregated data from the seminal SPRINT trial, which compared intensive versus standard blood pressure treatment goals on major adverse cardiovascular events.

**Results:** The method yields virtual populations, each with 9361 patients, faithfully mimicking the real SPRINT participants. Multiple logistic regression on 100 such populations shows that factors with the highest odds ratios for the primary event are, in descending order, past clinical cardiovascular disease, age ≥ 75, chronic kidney disease, high non-HDL, and smoking history. Intensive blood pressure treatment, the trial’s intervention, had an odds ratio of 0.74 [0.63-0.87]. On all these measures, the 100 RE-ViPPs mirrored the real SPRINT participants, including the intensive therapy result (actual SPRINT odds ratio: 0.74 [0.62-0.88]).

**Conclusions:** Clinical data dissemination has limitations. The most coveted data is descriptive at the individual level but comes with significant cost, effort, and time. There is potential for privacy breaches, and the open-data movement has progressed slowly due to data-ownership concerns. RE-ViPPs closely matched the true SPRINT population. Applied to trials, registries, and databases, RE-ViPPs could reduce open-data burdens by encouraging dissemination of aggregate cross-tabulated real data that allow investigators to generate and measure virtual patients.

## Introduction

Patient-level data from clinical trials, registries, and the electronic health record (EHR) is the basis for most clinical research. Obtaining and sharing such data can be challenging, and concerns have been raised about cost, effort, patient privacy, and data ownership [1]. Even journals with strong policies to promote data sharing of published papers have suboptimal adherence by authors [2]. Paying the right amount of attention to who requests individual level data, the purpose of the request, the structure and format of the data, and its de-identification to an extent sufficient to eliminate any chance of deducing identity, can be administratively taxing on the source institution, and often those costs are passed on to the requesting investigator [3–5].

Several entities have taken the initiative to open their data for specific trials and registries, which is a positive step forward. These include a consortium of clinical study sponsors initiating clinicalstudydatarequest.com [6,7], the Yale University Open Data Access (YODA) project [8], Duke’s Supporting Open Access Research (SOAR) initiative [9], ImmPort [10], and Project Data Sphere [11]. Some have associated costs in the thousands of dollars whereas others are free. Biolincc is the NHLBI’s free source for trial data and has reasonable access [12,13]. Besides clinical trials, sources of data can also include institutional, local, national, and international registries as well as clinical data warehouses driven by EHR data from one or more institutions. Typical time from request to obtaining individual-level de-identified data from one’s institutional EHR ranges from weeks to months and costs from hundreds to thousands of dollars [4,5].

However, there are sources of *aggregated* data, based on real clinical populations, which are currently underutilized for their potential. Public systems such as Center for Disease Control’s WONDER [14], the Healthcare Cost and Utilization Project (HCUPnet) [15], and the Behavioral Risk Factor Surveillance System Web Enabled Analysis Tool [16] are examples of this sort of aggregate open data for vital statistics, registries, and survey results, respectively. As for EHR data, the Informatics for Integrating Biology & the Bedside platform (i2b2) and similar platforms are deployed by multiple hospital systems to query data in an aggregate fashion at those institutions; they are mostly promoted as tools for cohort identification for planning future requests for individual-level data, but they have potential to be more widely used for research purposes [17–19]. Beyond informing requests for individual level data, these sources of aggregate data, as long as they can be queried in a richly cross-tabulated fashion, could contain enough information to approximate real populations. If this can be demonstrated, it would allow investigators using open data to analyze clinical populations without infringing upon patient privacy or overly taxing current systems in place, and at the same time, allow investigators running trials or maintaining registries to continue to have ownership of their individual-level data.

Here a method is described for creating populations of virtual patients from cross-tabulated aggregate data. It is applied to the SPRINT trial, a study of 9361 patients randomized to intensive versus standard blood pressure control, for which the individual-level data summarized in the original publication [20] was recently ‘opened’ through NHLBI’s Biolincc and is thus known [21]. SPRINT was chosen because it is a landmark trial with a well-defined population, and most importantly, provides verified individual-level data that allows for comparison of the replicate engineered virtual patient populations (RE-ViPPs) generated from aggregate data with the actual SPRINT population.

## Methods

### Study data

This is a secondary analysis of SPRINT data, which is provided publicly, in de-identified fashion, by the National Heart, Lung, and Blood Institute (NHLBI) via the Biolincc data repository (https://biolincc.nhlbi.nih.gov/home/). The study was deemed exempt by the University of Chicago IRB and the data was obtained after a signed data use agreement.

In SPRINT, participants were > 50 years old with a systolic blood pressure (SBP) of 130 to 180 mm Hg and had an increased risk of cardiovascular events for at least one of the following four reasons: clinical or subclinical cardiovascular disease, chronic kidney disease (CKD), a 10-year risk of 15% or greater on the basis of the Framingham risk score, or ≥ 75 years old. Patients with diabetes or prior stroke were excluded. Participants were randomized to an SBP target of either less than 140 mm Hg (the standard-treatment group) or less than 120 mm Hg (the intensive-treatment group). The primary composite outcome was myocardial infarction, other acute coronary syndromes, stroke, heart failure, or death from cardiovascular causes, measured over an average of 3.3 years [20].

The SPRINT individual-level data were aggregated by first choosing 9 independent categorical factors, all of which are well-established as influencing cardiovascular outcomes. Of these, eight were categorical in the original data [sex, senior (age≥75), black race, current or former smoker (grouped together in the present study), SBP above the highest tertile at the start of the trial, CKD (eGFR < 60 ml/min/1.73m^2^), history of clinical cardiovascular disease, and intensive SBP treatment]. The last (high non-HDL) was determined by calculating the non-HDL for each patient and identifying those with non-HDL > 160 mg/dL. The dependent variable was the primary composite outcome, also categorical.

### Construction of Replicate Engineered Virtual Patient Population (RE-ViPP)

One hundred RE-ViPPs were created. The starting material for RE-ViPP construction consists of aggregate data cross-tabulated for every pair of factors to be analyzed. The cross-tabulation is created by partitioning the entire population into subgroups by the status of one independent factor and recording, for each subgroup, the prevalence of each of the remaining factors, along with the rate of the dependent variable (**Supplementary Figure S1**, Steps 1-2). For SPRINT, this is straightforward, since the individual-level data is known. For EHR, registry, and large trial databases, this can be done, without giving investigators access to the individual level data, as long as the database can be queried for two factors at a time. The results of all this partitioning will yield a cross-tabulation of factor prevalences and outcome rates for each subgroup (as in **Figure 1** for SPRINT).

**Figure 1.**
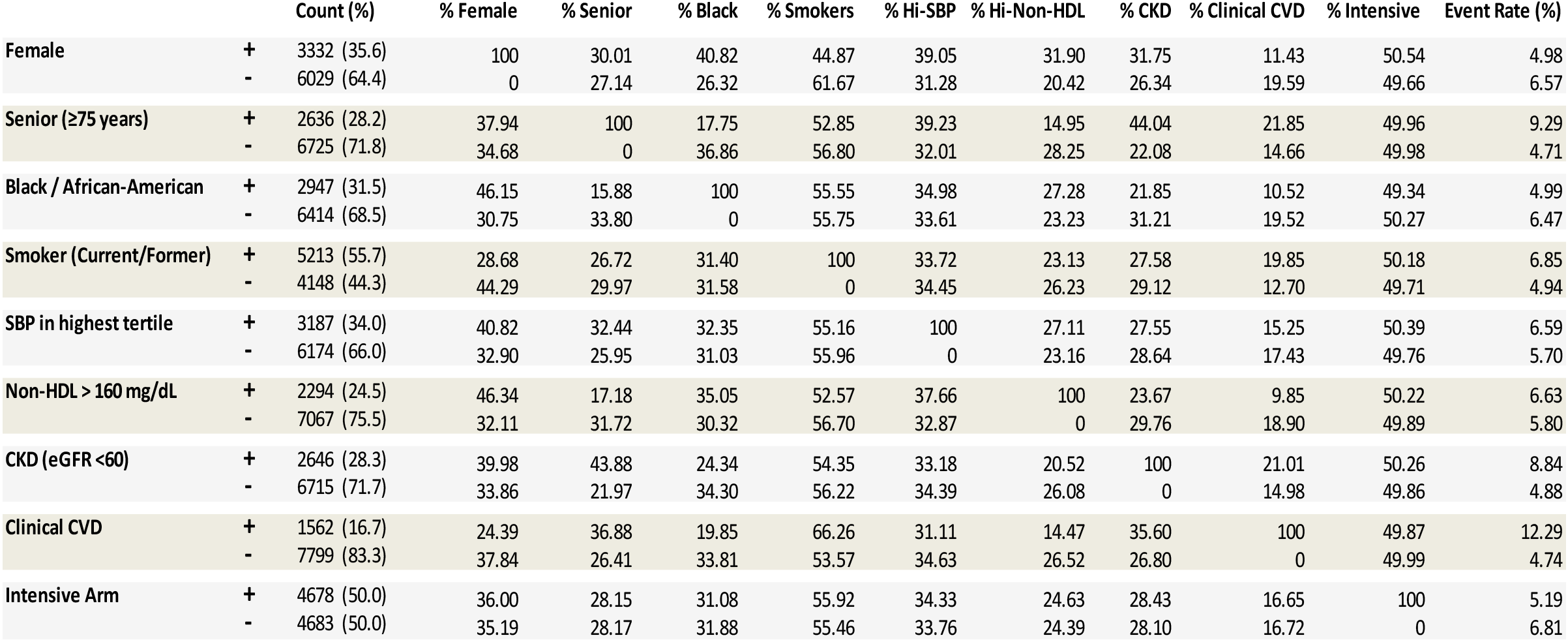
Cross-tabulated aggregate data from SPRINT. This represents the results from Steps 1-2 outlined in the methods and in Figure S1. The primary event rate and the prevalence of 9 independent factors are shown for subgroups defined by presence or absence of those same factors. The verbal equivalent of the top two rows, for instance, is: “Of the 3332 female participants of SPRINT, 100% were female, 30.01% were senior, 40.82% were black, [etc..], and 4.98% suffered the primary event. Of the 6029 non-female participants of SPRINT, 0% were female, 27.14% were senior, 26.32% were black, [etc..], and 6.57% suffered the primary event.” The data shown here is the only starting material required to generate RE-ViPPs.

With this cross-tabulation in hand, to start a RE-ViPP, the exact number of patients with and without the first independent variable is assigned to match the real population (**Supplementary Figure S1**, Step 3). In the case of the SPRINT analysis, this was, arbitrarily, female sex. Then, in those virtual patients with the first factor, the second factor (in SPRINT, this was senior status) was assigned to the exact percentage measured in the real population and the same was done for those virtual patients without the first factor. Thus the status of the first and second chosen factors will match the real population exactly.

Starting with the third factor (**Supplementary Figure S1**, Step 4), linear regression was performed on the aggregate data using all the previously assigned factors to estimate the chances of a patient having the new factor as a function of the previously assigned factors. The β coefficients and y-intercept from this linear regression comprise a formula to calculate a probability of having the new factor for each virtual patient (Supplementary Materials, Appendix 1). Significant interactions between pairs of factors are checked in the linear regression and included if the interaction β is within a factor of 1000 from the other β coefficients (in SPRINT, no such interactions were found between any pair of the 9 factors).

Once the probability (between 0 and 1) of having the new factor is calculated for each virtual patient, a random number (between 0 and 1) is generated for each patient (**Supplementary Figure S1**, Step 5). If the random number is less than the patient’s probability of having the new factor, the patient is assigned as having the new factor. Otherwise, the virtual patient does not. This repeated for each patient. Once all patients are assigned, the process repeats for the next factor to be assigned. After sequentially assigning the status for all the factors for all the virtual patients in the above manner, the outcomes are assigned in the same manner.

Once all factors and outcomes have been assigned to all virtual patients, the overall rate of each factor and outcome is measured against the actual population, and the virtual population is only accepted if within a specified tolerance, for all factors, of the actual population. If the rate of any factor is outside this tolerance, the random number generation is repeated for all patients and factors in order to create a new virtual population until a population is found that is within tolerance for all factors (**Supplementary Figure S1**, Step 6). For this study of SPRINT, the tolerance was set at ±0.25%. One hundred such acceptable RE-ViPPs are generated in the above manner.

### Analysis of RE-ViPPs

Each of the 100 RE-ViPPs is checked for correlation between the independent variables. It is then subjected to multiple logistic regression using the nine categorical independent variables and the primary composite outcome as the dependent variable. Random number generation, assignment of virtual patient factors, and tolerance testing were conducted in Microsoft Excel with Macros. The linear and logistic regression analyses were all conducted using R. Figures were created with ggplot2 in R.

### Patient and public involvement

Patients were integral to the SPRINT trial, and patient representatives were involved in the decision to make the trial data open through the National Heart, Lung, and Blood Institute. Patients were not involved in the design or implementation of the RE-ViPPs method or in the interpretation of the results, and results of this study cannot be disseminated directly to study participants due to the virtual nature of the data. The findings will be disseminated to the general population through national and international press, with the hope that it will encourage aggregate open data sharing.

## Results

The process for creating RE-ViPPs is presented in the methods section and summarized in **Supplementary Figure S1**. The process starts with curating cross-tabulated aggregate data. For SPRINT, this was obtained by analyzing the individual-level data. However, in its application for populations where individual-level data is not desired or easily available, the creation of the cross-tabulated data would come from querying an EHR database (such as via i2b2) or open aggregate/summary data from registries or clinical trials. **Figure 1** depicts the cross-tabulated aggregate data from SPRINT.

In the case of RE-ViPPs based on SPRINT, the populations were created based on 9 independent factors and the combined primary outcome event. Each RE-ViPP is composed of 9361 patients, utilizes eight equations (Supplementary Materials, Appendix 1), derived from linear regression analysis, for estimating the chance of a patient having a factor or outcome, and requires a total of 74,888 random numbers (9361 virtual patients × 8 random numbers per virtual patient). The tolerance was set at 0.25%, meaning that the prevalence of each factor and the primary event rate all had to be within this absolute percentage from the true rate in SPRINT. If this was not true for a given virtual population, all 9361 virtual patients were regenerated until the criterion was met. The “SPRINT RE-ViPP Generator” is available for download (Supplementary Materials).

One hundred RE-ViPPs were created in this randomized fashion, constrained by the chosen tolerance (each of these RE-ViPPs is available for download in Supplementary Materials). The rate of each included demographic factor (sex, age≥75, black race), clinical factor (prior/current smoking, SBP above the highest tertile, non-HDL>160 mg/dL, CKD defined by eGFR>60, past history of clinical CVD), trial factor (intensive vs standard BP therapy), and the primary outcome (composite myocardial infarction, other acute coronary syndromes, stroke, heart failure, or cardiovascular death), as required by the design, all adhere very closely to the SPRINT rates (**Figure 2**).

**Figure 2.**
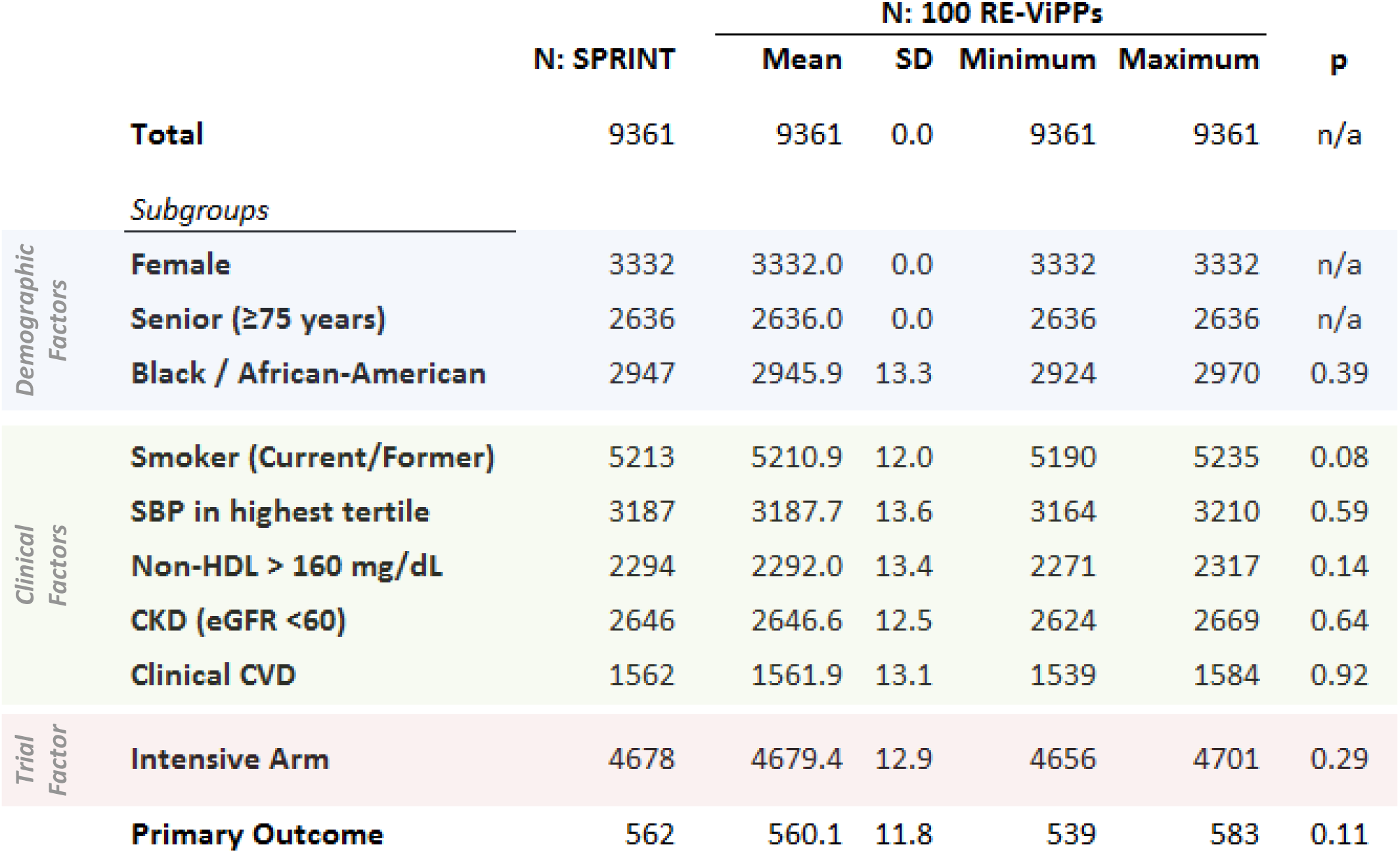
Patient counts in each RE-ViPP subgroup by demographic factor, clinical factor, trial factor, and primary outcome all adhere closely to SPRINT. One hundred replicate populations, each with 9361 virtual patients, were generated. The rates were, by design, within an absolute 0.25% of the actual SPRINT trial for every factor in every RE-ViPP. The full range of RE-ViPP counts are shown, as are p values from one-sample t-tests comparing the 100 RE-ViPPs to the actual SPRINT participants.

Also across all subgroups, the prevalence of each demographic and clinical factor in each of the 100 virtual patient populations clusters around the actual rates in SPRINT (**Figure 3**). The virtual patient population is randomized well to intensive and standard BP therapy, just as the real SPRINT patients, and primary event rates also match across all subgroups (**Figure 3**). It is worth noting that the SPRINT population has higher female and lower senior prevalence amongst black participants, lower rates of smoking and higher non-HDL amongst women, more frequent CKD amongst seniors, and higher event rates in seniors, those with CKD, those with past clinical CVD, and those randomized to non-intensive treatment. All these features are preserved in all of the virtual patient populations.

**Figure 3.**
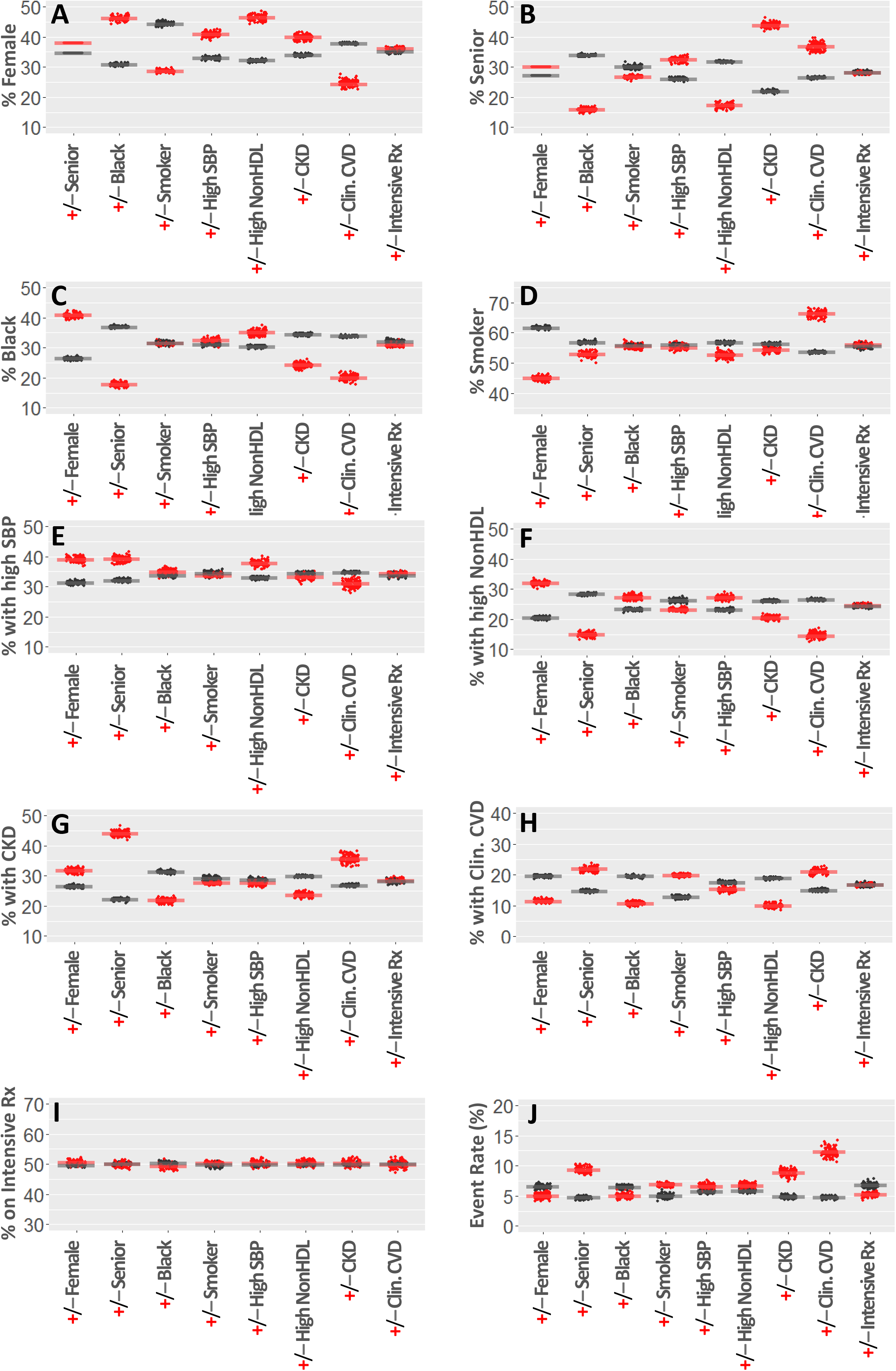
RE-ViPPs match the real SPRINT population. In each plot, a dot represents a single virtual patient population subgroup and horizontal bars are the real SPRINT rates. Red and gray respectively depict the subgroups with and without the corresponding x-axis factor. The prevalence of demographic factors [female sex (A), age≥75 (B), and black race (C)] and baseline clinical factors [prior/current smoking (D), highest tertile SBP (E), non-HDL>160 mg/dL (F), CKD defined by eGFR>60 (G), and past history of clinical CVD (H)] in each of the 100 virtual patient populations clusters around the actual rate in SPRINT across all subgroups. The virtual patient population is randomized well to intensive vs. standard BP therapy (I) across subgroups just as the real SPRINT patients, and primary event rates also match across all subgroups (J). Notable features of the SPRINT population include higher female and lower senior rates amongst black participants (A-C), lower rates of smoking and higher non-HDL amongst women (D,F), higher CKD amongst seniors (G), and higher event rates in seniors, those with CKD, those with past clinical CVD, and those randomized to non-intensive treatment (J). All these features are preserved in all of the virtual patient populations.

Beyond subgroup characteristics, the RE-ViPPs can be analyzed on the individual level similar to a multivariate analysis performed on real populations. Correlation matrices show no strong correlations between the analyzed factors in the RE-ViPPs, just as in the real SPRINT population (**Supplementary Figure S2**).

Multiple logistic regression performed on each of the 100 virtual populations show that the odds ratios (ORs) for each factor closely approximates the ORs in SPRINT (**Figure 4**). The confidence intervals of the RE-ViPP ORs are also similar to SPRINT. A volcano plot shows the OR plotted against p values for each factor in each RE-ViPP, again demonstrating close adherence to SPRINT in terms of both magnitude and significance of the findings (**Figure 5**). The risk factors with the highest odds ratio for the primary event are, in descending order, past clinical cardiovascular disease, age ≥ 75, chronic kidney disease, high non-HDL, and smoking history. Factors associated with fewer events are female sex and intensive blood pressure treatment. Just as with the true SPRINT data, neither black race nor high initial SBP had confidence intervals that clear the line of neutrality.

**Figure 4.**
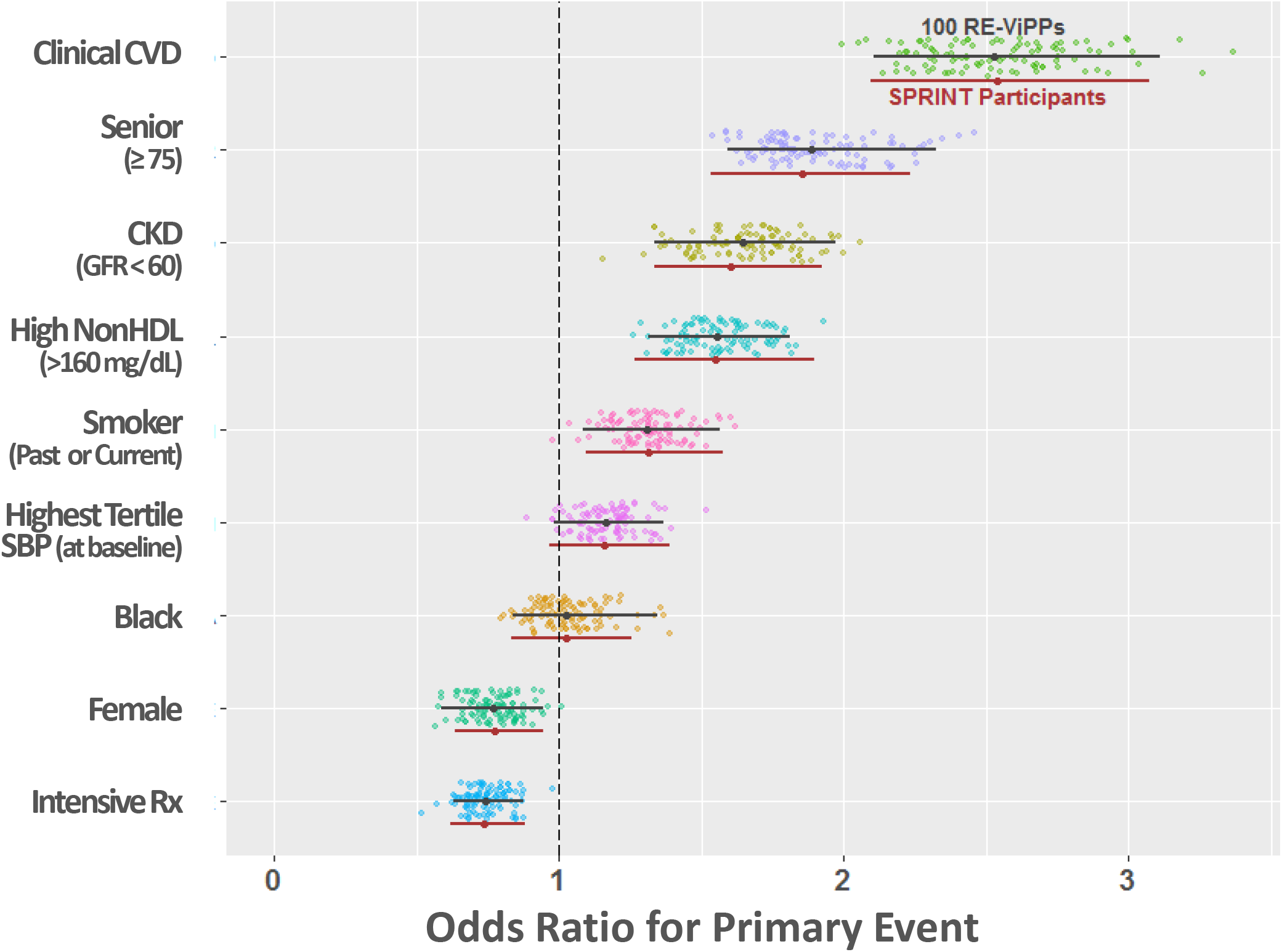
Multiple logistic regression of RE-ViPPs. Multiple logistic regression performed on the 100 virtual populations (9361 patients each) show that the Odds Ratio (OR) for each factor approximate those of SPRINT. Each colored dot is the calculated OR for a virtual patient population. The OR calculated from the mean beta coefficient for each factor along with the central 95% of these ORs is shown in black. For comparison, the actual SPRINT OR and 95% confidence interval for each factor is shown in red.

**Figure 5.**
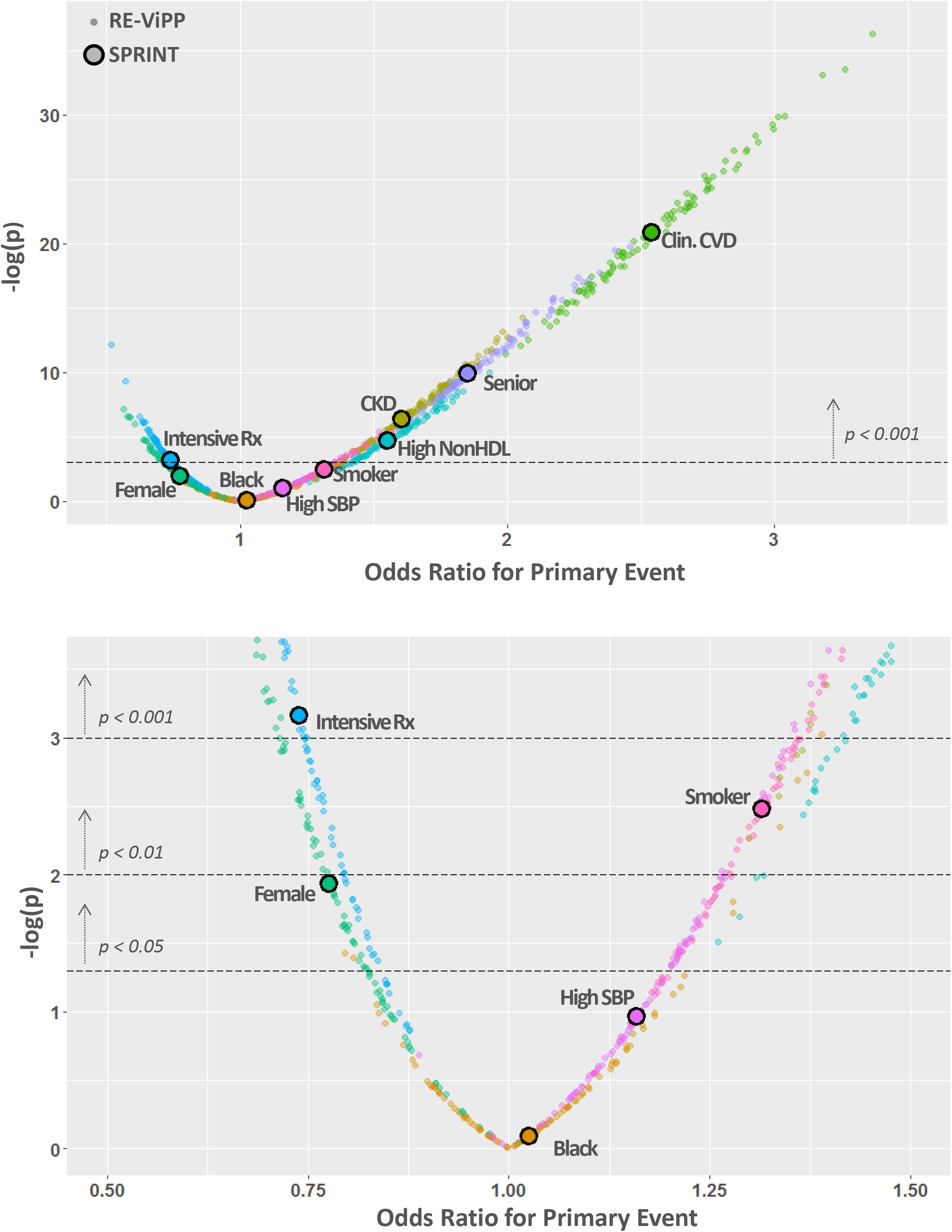
Multiple logistic regression of RE-ViPPs match SPRINT. Volcano plot shows the OR plotted against the −log of the p value for each factor (color-coded) in each RE-ViPP and in SPRINT. Lower panel is a magnified area from upper panel. The magnitude and significance of the each factor’s impact in SPRINT is recapitulated the by RE-ViPPS, as the SPRINT value falls in the middle of the values for the RE-ViPPs for each respective factor.

## Discussion

The approach described here was chosen to demonstrate that meaningful information can be gleaned from certain aggregated data to simulate individual patients. The method extends beyond what is traditionally done with summary information and it can approximate the results from the true individual-level data. It is also distinct from prior methods that rely on individual-level data to create virtual patients for the purposes of physiological experiment simulation or clinical trial simulation [22,23]. It is very important to emphasize that the method cannot be done with any form of aggregate data—the source data must be cross-tabulated by the studied factors and outcomes, and, as such, all the factors should be categorical, either nominal or ordinal. In the current example with SPRINT, the variables were binary, but this process could be extended to incorporate a limited number of non-binary categorical variables, in which case the cross-tabulation should account for all values of those variables. Another important consideration is the quality of the source data. In a clinical trial such as SPRINT, it is likely very reliable, but in EHR databases, variables such as race and smoking status can be variably and inconsistently reported, and diagnoses and outcomes may rely on ICD codes which primarily serve a billing/documentation function and may not be subject to rigorous independent verification [24,25]. Another limitation is that although the method checks and includes any pairwise interaction between variables that is large and significant, it does not address 3-way or more complex interactions. There were no important interactions between the variables chosen for this SPRINT study, and for other studies this limitation could be addressed by checking for such complex interactions during the linear regression analyses and/or by choosing, when possible, sufficiently distinct variables to analyze.

The method is reminiscent of imputation, the process by which missing data for a variable is replaced by a reasonable prediction, often using regression models, based on the extant values for that variable as a function of the other variables in the dataset [26,27]. This has perhaps taken its most robust form in multiple imputation of chained equations [28,29]. The method here is distinct from multiple imputation in several ways, but an analogy would be that every patient’s status for all the variables beyond the first two are treated as missing, and they are imputed sequentially based on logistic regression analysis of the percentage distributions in the aggregate data.

The current study is meant to serve as a practical proof-of-principle example. Data known at the individual level was used to highlight the method and show that the RE-ViPPs result matches that of the real population. In practice, this method would be most useful in situations when the individual-level data is either impossible or difficult to obtain. These are common situations. In many cases, there are concerns about data ownership or patient privacy [30]. In others, it can also involve excess time, effort, and cost [3,31]. Still in others, the issue may be one of data standardization, where a study would ideally be done across datasets or platforms, but the data exist in incongruent formats in each platform [32]. Also, the sheer bulk of individual-level data may be prohibitive, particularly when there is a desire to combine data from multiple EHRs across the world. With these applications in mind, the “SPRINT RE-ViPP Generator” is provided in the Supplementary Materials and can be customized to other sets of aggregate data and variables.

There are many recent high profile examples, not only in healthcare, of bulk individual-level data being hacked or leaked with major fallout [33,34]. In all these situations, keeping the individual level data undisclosed while providing the requisite summary statistics could allow other investigators to study the data while abrogating these concerns. One way this could be done is to keep the individual level data behind a firewall but allow conditional queries to extract summary data – this is what the i2b2 platform and others do for EHR data warehouses, what Center for Disease Control’s WONDER provides for vital statistics and other limited clinical information, and what the Behavioral Risk Factor Surveillance System Web Enabled Analysis Tool does for its large national health survey results [14,16,17,19]. Another approach would be to have the “data owners” provide the richly cross-tabulated results across as many categorical variables as possible as a small static file (like **Figure 1**) and make it open to interested investigators; this approach would probably be more feasible and require less digital infrastructure for individual clinical trial results in which full data releases are not planned. This could also allay concerns about the provenance of the source data in secondary analysis, as the shared information would be more succinct and verifiable. In all the above scenarios, it is worth emphasizing that none of the analyzed individual patients derived from this method are real. Patients in the real population are well represented by the virtual patients, but as digital versions of creation, the virtual patients are nameless composites only [35], with no opportunity to deduce identity.

This method should allow us to analyze trial, registry, and routinely collected clinical information over large populations to find potential connections between heterogeneous clinical and demographic factors in a novel way that does not directly access patient-level data. The method could reduce barriers that currently impede access and use of open data. It would thereby stimulate more meaningful use of extant information, ultimately to generate new hypotheses for further investigation in real individuals and to identify previously unrecognized clinical relationships.

## Author Contributions

Dr. Alenghat had full access to the aggregate data, preformed the analysis, and takes responsibility for its accuracy

## Data sharing statement

Access to SPRINT data is available from the National Heart, Lung and Blood Institute via Biolinnc. Access to the 100 RE-ViPPs is provided in the Supplementary Materials.

