## Supplemental Figures for "Replicate Engineered Virtual Patient Populations as Surrogates for Real Patient-Level Data"

### Figure S1

1 Partition entire real population of N Individuals with M categorical factors by whether they have Factor $X_i$

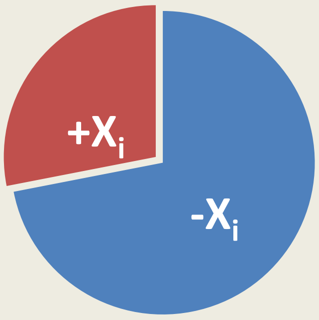


2 Measure prevalence ($P_i$) of all factors as well as outcome rate (Z) for those $+X_i$ and $-X_i$

| $+X_i$ | $-X_i$ |
|---|---|
| $P_1$% with Factor $X_1$ | $P_1$% with Factor $X_1$ |
| $P_2$% with Factor $X_2$ | $P_2$% with Factor $X_2$ |
| . | . |
| . | . |
| $P_M$% with Factor $X_M$ | $P_M$% with Factor $X_M$ |
| Z% with Outcome | Z% with Outcome |

Repeat 1 and 2 for each factor $X_1..X_M$

3 Start a Virtual Patient Population of N individuals. Assign the exact number of patients with and without Factor $X_1$ to match the real population. Of those with Factor $X_1$, assign the exact $P_2$% to have Factor $X_2$ and the rest lack Factor $X_2$. Of those without Factor $X_1$, assign the exact $P_2$% to have Factor $X_2$ and the rest lack Factor $X_2$. For Factors $X_1$ & $X_2$, the virtual population will match the real population exactly.

4 Starting with Factor $X_{i=3}$ and proceeding sequentially through Factor $X_M$, perform linear regression using $P_1..P_{i-1}$ & $P_1..P_{i-1}$ for each Factor $X_1..X_{i-1}$ as independent variables and $P_i$ & $P_i$ for $X_1..X_{i-1}$ as the dependent variable to determine coefficients $\beta_1...\beta_{i-1}$ and intercept Y. Check and include significant interactions between pairs of factors. Use $\beta_1...\beta_{i-1}$ and Y and already assigned $X_1$-$X_{i-1}$ in the virtual population to determine the probability, $PROB_i$ of having $X_i$ for each virtual patient. Repeat for the outcomes Z and Z using $P_1..P_M$ and $P_1..P_M$ for all Factors as the independent variables.

5 Generate a random number, $R_i$, between 0 and 1 for each factor $X_i$ (starting at i=3) for each virtual patient. If $R_i < PROB_i$, then assign the patient to have $X_i$. Otherwise, assign the patient to lack $X_i$. Do the same for $R_z$ and $PROB_z$ assigning the outcome to each patient.

6 Repeat 5 by regenerating the N•(M-1) random numbers until the prevalence of each Factor $X_1$-$X_n$ and the outcome rate are all within an tolerance T of the entire real population. Once this occurs, generation of a single virtual patient population generation is complete. Repeat 5 & 6 to generate additional replicates until the desired number of virtual populations are obtained.

**Figure S1. Replicate Engineered Virtual Patient Populations (RE-ViPPs): A step-by-step method.** The source material can be created by querying a database of accessible clinical data (Steps 1-2) or be provided by the data owners. Generation of virtual population (Steps 3-6) relies on assigning variable values to each virtual patient in a sequential manner based on linear regression models, driven by random number generation, and subjected to tolerance limits. Steps 5-6 are repeated to make replicates of the virtual popluation (in this case, 100 RE-ViPPs were generated for SPRINT).

### Figure S2

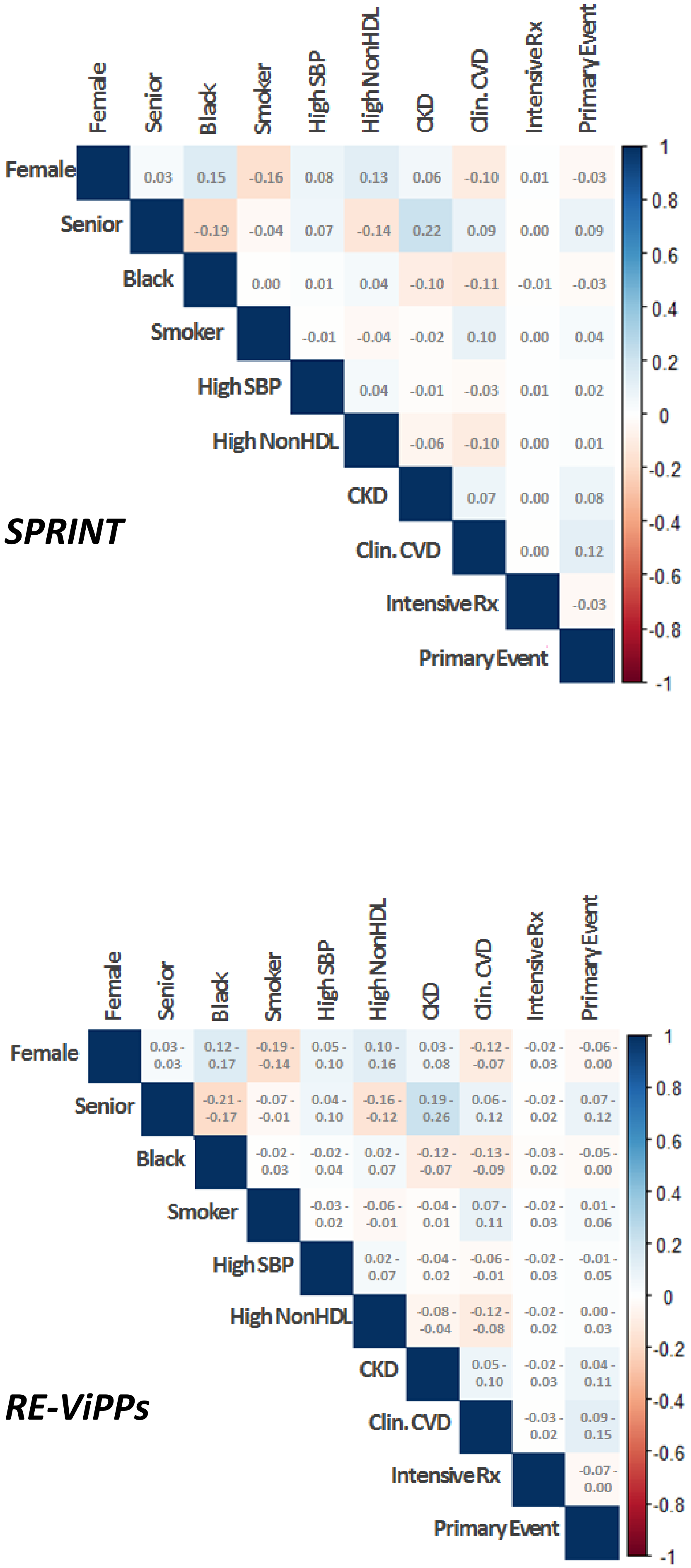


**Figure S2. Correlation coefficients of SPRINT and RE-ViPPs.** Correlation coefficients are shown for each pair of factors in SPRINT (upper panel). For the RE-ViPPS (lower panel), the average correlations for each pair of factors is reflected by the color scale, and the full range of correlations is provided. The pattern is similar to SPRINT. No coefficient exceeds 0.3 in any population.
